# AN ARTIFICIAL INTELLIGENCE FOR RAPID IN-LINE LABEL-FREE HUMAN PLURIPOTENT STEM CELL COUNTING AND QUALITY ASSESSMENT

**DOI:** 10.1101/2023.03.20.533543

**Authors:** Britney Ragunton, Steve Van Buskirk, Devin Wakefield, Ninad Ranadive, Andrew Pipathsouk, Baikang Pei, Hong Zhou, Tracy Yamawaki, Mike Berke, Chi-Ming Li, Christopher Hale, Songli Wang, Stuart M. Chambers

**Affiliations:** Genome Analysis Unit, Amgen Research, South San Francisco, CA; Discovery Technologies, Amgen Research, Thousand Oaks, CA; San Jose State University, San Jose, CA

## Abstract

The current state-of-the-art in hPSC culture is a bespoke and user-dependent process limiting the scale and complexity of the experiments performed and introducing operator-to-operator and day-to-day variation. Artificial intelligence (AI) offers the speed and flexibility to bridge the gap between a human-dependent process and industrial-scale automation.

We evaluated an AI approach for counting exact cell numbers of undifferentiated human induced pluripotent stem cells in brightfield images for automating hPSC culture. The neural network generates a topological density map for accurate cell counts. We found that the image-based AI algorithm can determine a precise number of hPSCs and is superior to fluorescence-labeled object detection; the algorithm can ignore well edges, meniscus effects, and dust, achieving an average error of 5.6%. We have built a prototype capable of making a go/no go decision for stem cell passaging to perform 26,400 individual well-level counts from 422,400 images in 12 hours at low cost.

---

There is an increasing demand for this technology that is labor intensive and sensitive to user variance and genetic diversity of stem cell lines. Currently, the dynamic process of pluripotent stem cell derivation and culture demands a highly skilled individual with years of experience. Automated platforms for stem cell culture would accelerate access to hPSC technology for drug development and clinical application of hPSC derivatives. We set out to identify a machine learning algorithm to standardize stem cell cultures.

In current practice, measuring the density of cells in a dish requires the cells to be dissociated and counted or have their nuclei stained and imaged in the dish. These methods are time-consuming and result in loss of cells. It is also common practice to measure confluence with an Incucyte with brightfield imaging, but that is often insufficient for measuring hPSC density given the range of cell compaction and area. We sought to streamline this process by counting cells directly from brightfield images of the live cells using machine learning technology.

Machine learning is a field of artificial intelligence in which computers can learn a task such as pattern recognition, classification, and prediction from data without relying on a predetermined equation or model. For image analysis deep learning, a subset of machine learning, the computer learns to perform tasks directly from data. Deep learning uses convolutional neural networks (CNNs) for learning features, representations, and tasks directly from the images. This differs from conventional machine learning in which features for classification are manually defined. Machine learning has been applied successfully in biologic applications to predict fluorescent labels from images (Christiansen et al., 2018), to evaluate hPSC quality (Wakui et al., 2017), to identify differentiated stem cells without molecular labeling (Buggenthin et al., 2017; Kusumoto et al., 2018; Waisman et al., 2019), and to identify macrophage subsets (Rostam et al., 2017).

Crowd counting is an application of machine learning that uses detection, regression and density estimation-based methods to count people in images of crowds. CNN approaches have been developed to aid crowd counting, including a new method using a cascaded convolutional network that learns both crowd count classification and a density map estimation (Sindagi and Patel, 2017). This new method has lower error and higher quality density maps compared to previous methods (Sindagi and Patel, 2017).

We hypothesized that the deep learning crowd counting model could be applied to counting cells in images. The model makes use of CNN layers to identify key features in the images that allow for accurate and selective cell counting and the creation of a density map. The cell count is important for plating at specific densities for passaging and differentiations. In addition to a cell count, the model also creates density maps that contain more information that can be used to assess the quality of the cells. We created a proof-of-concept platform to monitor hPSC growth and make passaging decisions with our trained model by using a Precise Automation robotic arm, Cytomat automated incubator, Celigo Imaging Cytometer, and Overlord laboratory automation software. The method can be used across all hPSC cultures as is or by the addition of more data through transfer learning.

## A crowd counting convolutional neural network can be trained to detect and count individual hPSCs

We leveraged neural networks developed to generate accurate predictions of crowd counts to accurately predict the number of hPSCs within a colony by hypothesizing that hPSCs resemble heads in a crowd. A CNN shown to have superior performance in crowd counting (Sindagi and Patel, 2017) was used to generate both an accurate count and physical location of the cells by generating a three-dimensional density map in which density is distributed along x- and y-coordinates of the microscope. To test whether this CNN could be trained to detect hPSCs at single cell resolution, aligned bright field and Hoechst-stained images of hPSC colonies plated in 96-well format were obtained using an automated microscope (**Figure 1 A, B, C**). Centroids were identified using the Hoechst-stained images and converted to a density map using ImageJ/FIJI to serve as ground truth for training the AI (**Figure 1D**). Additional example images can be found in **Supplemental Figure 1**.

**Figure 1.**
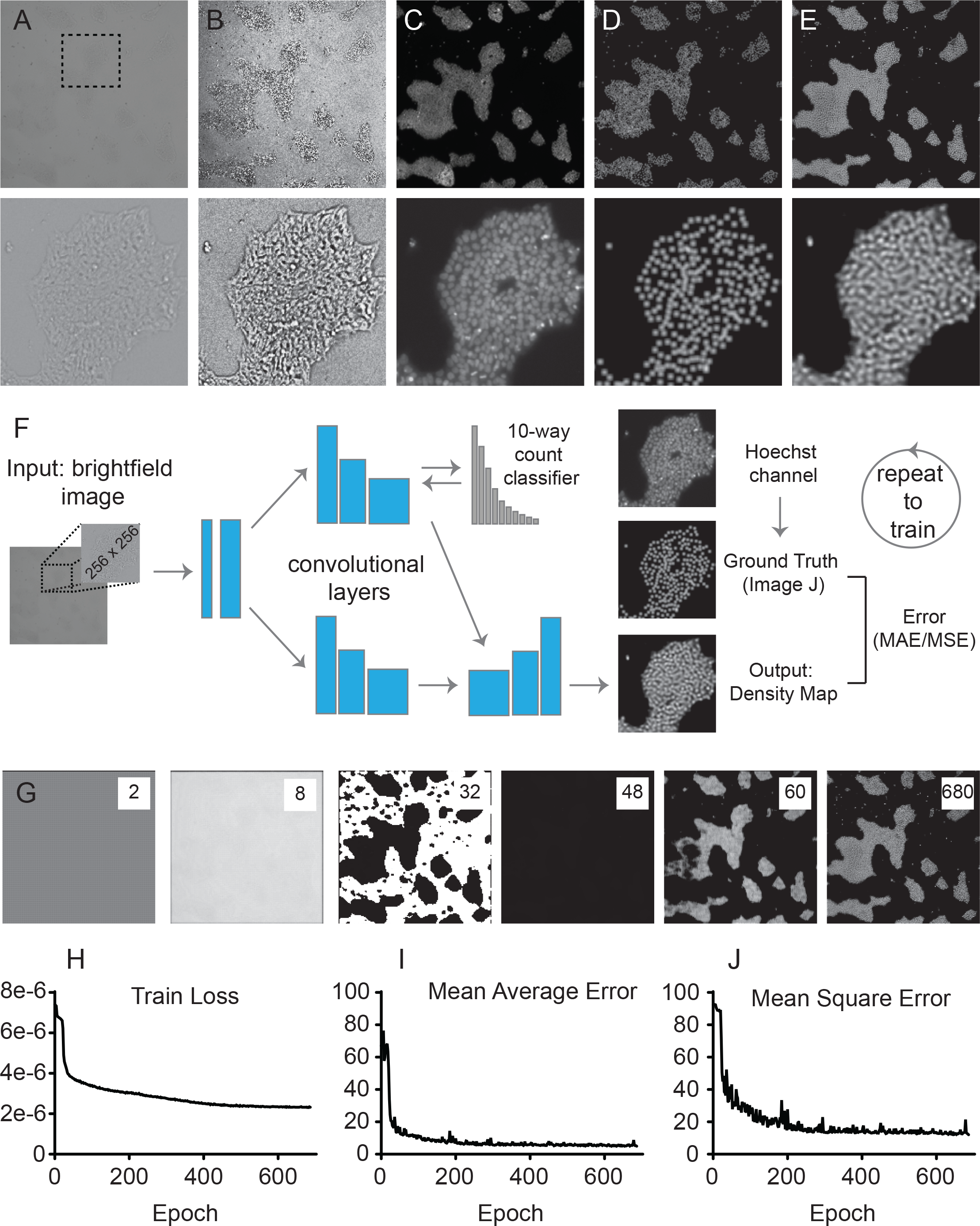
Model training workflow. A) Representative image from the dataset used for developing the cell counting model. The top row of images show the whole field of view and the second row of images are a crop shown by the box in the first image. The first column of images are brightfield. B) Enhanced version of the brightfield image. C) Hoechst stained image. D) Ground truth image derived from the Hoechst image. E) Density map output by the model. F) Structure of the deep learning model used. G) The density map output from the model of an example image as training progresses. Each image is labeled by the epoch. H) The cross-entropy loss of the training set over the training epochs. I) The mean average error (MAE) of the validation set over the training epochs. J) The mean square error (MSE) of the validation set over the training epochs.

The image-based CNN is comprised of two parallel processes filtered through convolutional layers (**Figure 1F**). One half (top) approximates counts and bins an image into a 10-way count classifier with the intention of binning images based on the approximate number of cells within the field. The information is used to inform the second half (bottom) to generate a density map with local maximal densities representing individual nuclei. Through repeated training rounds (epochs) the connections between individual layers of the CNN are either strengthened or weakened on the basis of similarity of the model-generated density map to the ground truth density map. Images were run through intermediate models captured at even epochs and the results were output as density maps to capture the CNN training process. Interestingly, the AI underwent a trial-and-error phase from epoch 8-48 before determining the proper density localization, including an inverted density map in which density was assigned to the empty portions of the dish before finding the correct approximate distribution (**Figure 1G**). By epoch 60, gross colony morphology and positions were correctly determined and further refined throughout iteration. This learning process coincided with a minimization of the train loss, mean average error, and mean square error (**Figure 1H**). The training was halted to select the model from epoch 680 as a minimum mean average error and mean squared error. The resulting model was found to ignore microscopic artifacts including particulate on the bottom of the microwell plate, well edges, bubbles, and variation in focal planes, without the need for augmentation of the original bright field images (**Supplemental Figure 1**), and was used in subsequent experiments. We are augmenting the model for the more challenging task of analyzing phase-contrast images, which will allow historical analysis of published micrographs.

## Model performance and implementation of the cell counting CNN on an automated platform

The model we developed is data-rich; it can localize the relative positions of cells in the dish, be sub-divided to generate counts within specific fields of view and summarize larger areas by calculating the area under the curve. To illustrate this point, images can be sliced at given horizontal coordinates and the gray value plotted. Comparison of the magnitude and sharpness in features of the low-contrast original images and the density maps revealed by the AI illustrates the information transformation performed by the trained AI model (**Figure 2A**). The model was evaluated using newly imaged data as a correlation of the ground truth, provided by fluorescence based object detection, to the model results demonstrating a R-squared value of 0.994 (**Figure 2B**).

**Figure 2.**
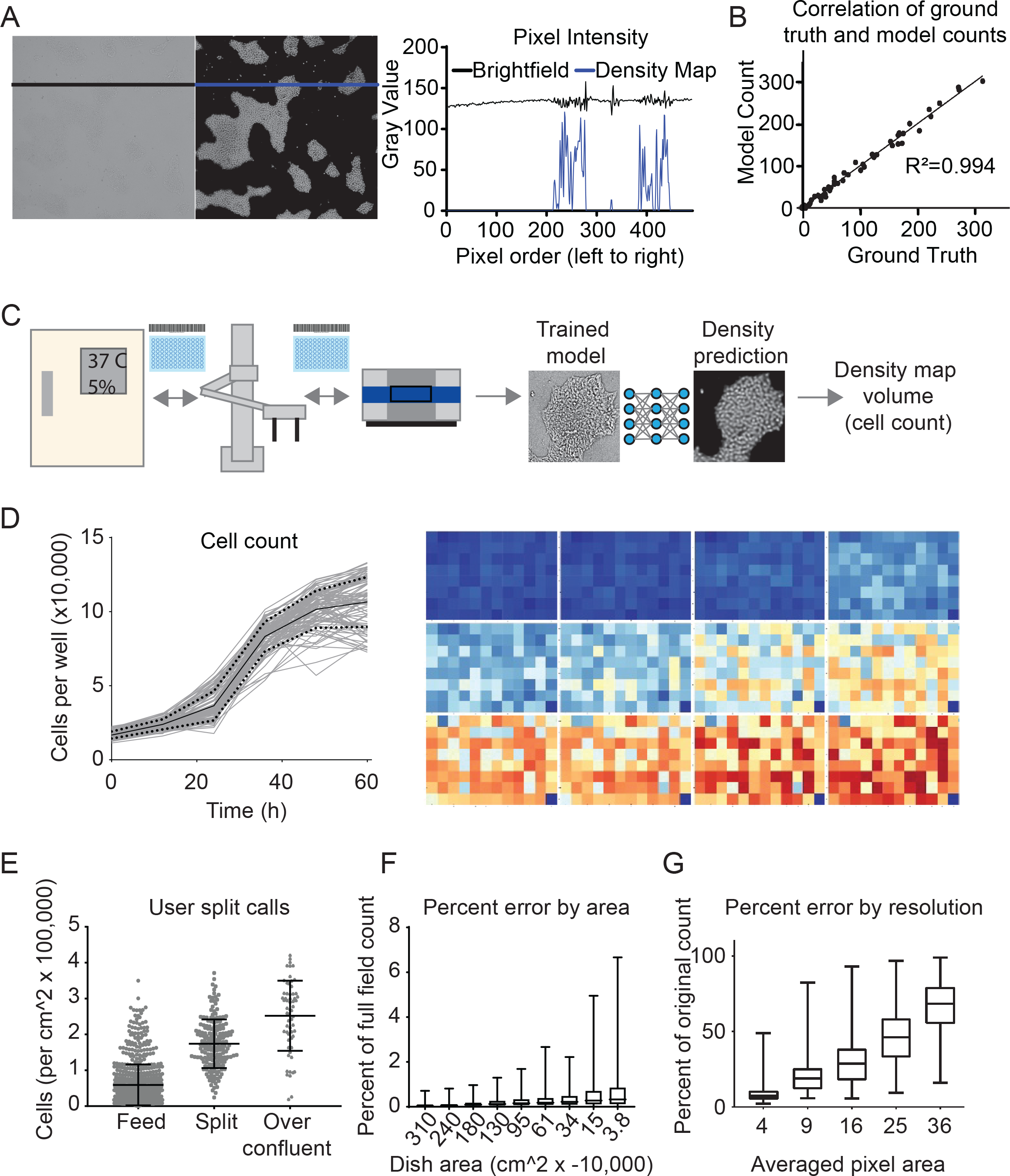
Integration of the AI Model into Automated System. A) Plot showing the difference in pixel intensity at the line drawn across the images for brightfield and ground truth. B) Correlation between ground truth determined by image object detection and the AI model on data not used for the deep learning. C) An automated platform was created to monitor hPSC growth by integrating a Cytomat automated incubator (Thermo Scientific), robotic arm (Precise Automation), Celigo imaging cytometer (Nexcelom Bioscience) and Overlord laboratory automation and control Software (PAA). After imaging, images are automatically sent to an AWS S3 bucket where they are run through the cell counting model. D) Model output from automated runs to confluency. Plate wide heat maps and growth curves show the change in cell density across a plate as the cells grow to confluency. E) Split decision training was performed with data from the automated platform. Plot of split decisions based on cell counts per unit area. F) Plot of percent error from area down-sampling. G) Plot of percent error from resolution down-sampling.

We next built an automated system consisting of an automated cell incubator, robotic arm, and the identical automated microscope used for acquiring the training data (**Figure 2C**). Full-well images from 27 bar-coded 96-well tissue culture plates were recorded every 12 hours. Images were automatically uploaded to the cloud and cell counts and heat maps were calculated to monitor cell growth over the time course (**Figure 2D**), demonstrating the in-line performance of an automated system over the time of hPSC growth.

Add some discussion here about figure 2 E, F & G:

Split decision training was performed by classifying images of hPSCs that should be fed, split or are considered over confluent (Figure 2E). A power analysis was also performed to see if using a smaller area or lower resolution gave comparable results (Figure 2 F, G).

## Methods for assessing quality of induced pluripotent stem cells

We asked whether the same bright field data could be used to determine culture quality, specifically whether machine learning could detect differences in cell morphology due to differentiation. Six additional induced pluripotent stem cell lines were generated using the same methods. The lines, named C2 through C7, were selected for their varying degree of apparently differentiated cells observed upon continued passaging. PluriTest (Muller et al., 2011), an unbiased bioinformatic method for accurately determining the pluripotency of human stem cells (Initiative, 2018) was used to establish a quality score and ranking for the individual lines. Bulk mRNA sequencing was performed on all seven lines, resulting in a PluriTest ranking from best to worst: LT, C7, C3, C4, C5, C6, C2 (**Table 1** and **Supplemental Table 1**). The percentage of cells triple positive for pluripotency factors SSEA-3, SSEA-4, and TRA1-60 measured by flow cytometry also correlated well with this ranking (**Supplemental Figure 2**).

**Table 1.** Ranking quality of hPSC lines using different methods. Standard methods for evaluating quality including PluriTest and gene expression were used as well as new deep learning approaches.

| PluriTest | Manual Ranking | Rank by FACS | Gene Expression | Image Classification | Semantic Segmentation |
| --- | --- | --- | --- | --- | --- |
| C2 | C2 | C2 | C2 | C2 | C2 |
| C6 | C4 | C6 | C4 | C4 | C6 |
| C5 | C6 | C5 | C6 | C6 | C4 |
| C4 | C5 | C3 | C5 | C5 | C5 |
| C3 | C3 | C4 | C7 | C3 | LT |
| C7 | C7 | LT | C3 | LT | C3 |
| LT | LT | C7 | LT | C7 | C7 |

Density maps were generated for the new hPSC lines using the same cell counting model, demonstrating in comparison to Hoechst stained images, the accuracy and broad applicability of the model to additional hPSC lines (**Figure 3A**). A normal q-q plot of the cell counts for the hPSC lines was used determining how closely the sampling of individual fields of view fit a normal distribution. As an example, if differentiation of cells led to confluency across all fields of view, as is seen for C2, the result was a ‘middling effect’ in which images with median counts are observed more frequently, seen as a concave down curve. The clones exhibit a similar rank order by plot curvature and deviation from the normal distribution (**Figure 3B**).

**Figure 3.**
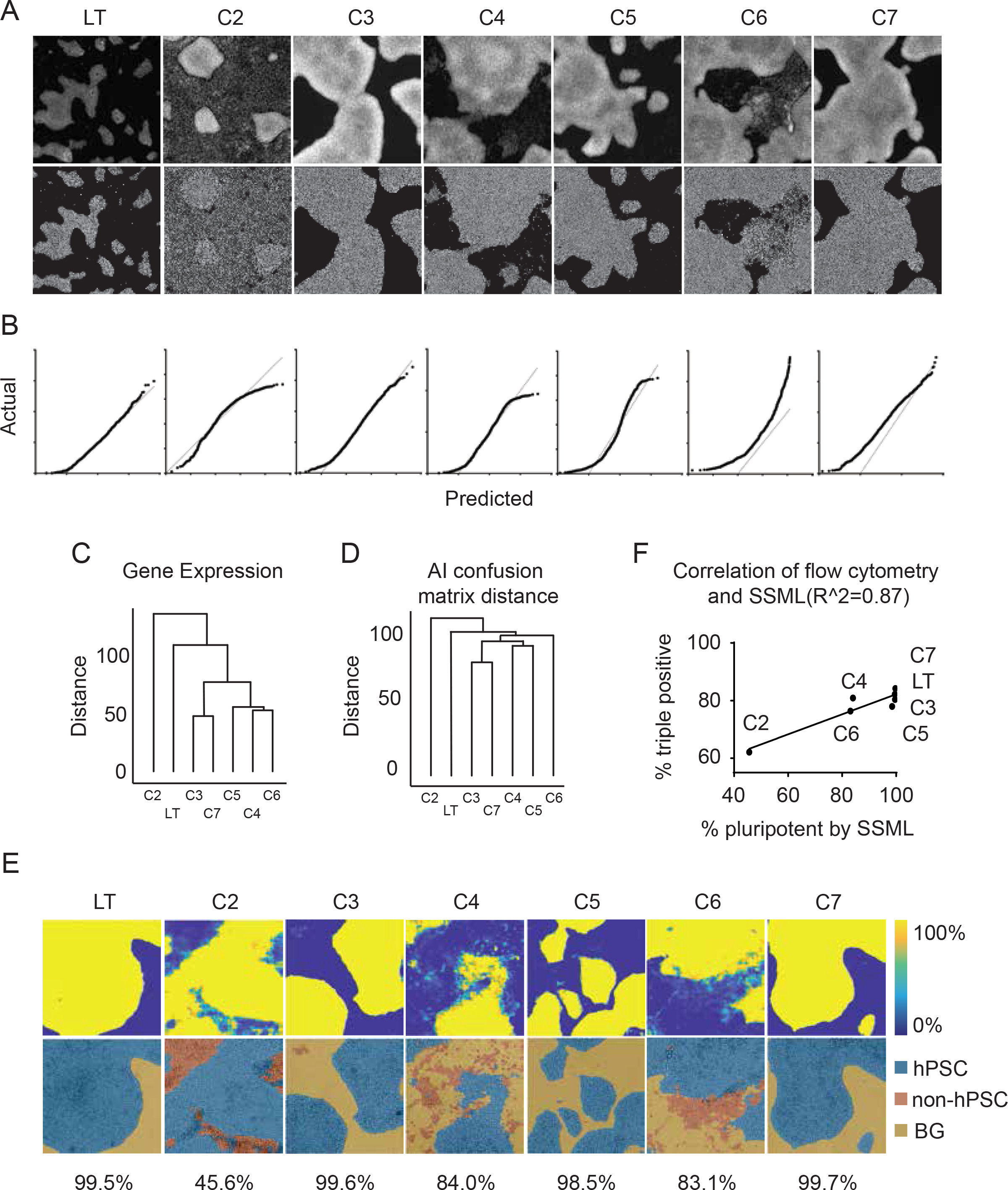
Determining hPSC Quality. A) Nuclear stained images from each hPSC clone and the corresponding density map outputs. B) Normal qq plots for each hPSC clone. C) Dendrogram created from RNASeq data using top 500 genes, Euclidian distance, and complete linkage D) Dendrogram created from a confusion matrix generated when evaluating performance of the deep learning model for clone identification using Euclidian distance and the single-link agglomerative method. E) The top row of images are hPSC pixel score maps from each hPSC clone. The second row of images show the segmentation of pixels in images from each hPSC clone. Blue is hPSC pixels, orange is non-hPSC pixels, and yellow is background (BG). Percentage of hPSC pixels from the total cell pixels for each clone. F) Correlation of flow cytometry and SSML.

Using this collection of clones, pre-trained networks available through MATLAB (Mathworks) were evaluated and selected for the best accuracy in conducting three AI tasks for determining clone quality: distinguishing between clones, classifying images as undifferentiated or differentiated, and semantic segmentation. We hypothesized that an AI may be able to distinguish between clones from a single image, but may struggle with highly similar clones, and the presence of differentiated cells could serve as one distinguishing characteristic. In support of this hypothesis, the highest accuracy for this method amongst the four models tested was 79.36% (**Supplemental Table 2**), lower than one would expect if the AI could perfectly distinguish clones. However, the resulting confusion matrix could be used to generate a dendrogram on the basis of distance between clones, which appears strikingly similar to the gene expression dendrogram for the clones (**Figure 3C, D**), supporting the notion that simple brightfield images contain sufficient information for evaluating hPSCs.

For image classification, the training data were generated by binning individual images for each clone into either pluripotent or differentiated classifications until a balanced data set of similarly sized bins was created for each of the clones. The AI was then trained to distinguish between images that showed undifferentiated hPSCs and one that contained differentiated cell types, achieving a 95.15% accuracy. This model was then used to evaluate and rank each clone based on the frequency of encountering images containing differentiated cells (**Table 1, Supplemental Table 3**).

Semantic segmentation is an AI method of classifying individual pixels in images such that an AI can make predictions and report a percent likelihood that each pixel of an image falls into a given class. Thirty-two random images were selected for each clone and pixels were user-painted according to the three classes –undifferentiated hPSC, differentiated cell, and background. After training, the final model had an accuracy of 95.99%. The AI results can be visualized as either a percent likelihood for each class or a combined pixel painted image (**Figure 3E)**. Frequency of hPSC pixels to total pixels that contained cells were calculated to score and rank the clones according to their pluripotency. The frequency of undifferentiated hPSC pixels strongly correlated with the percentage of triple positive cells measured by flow cytometry (**Figure 3F**), indicating that the semantic segmentation AI can successfully estimate the cellular composition for each hPSC clone and report a quantitative score that can be used to rank the clones (**Table 1**).

Cell therapies using derivatives of human pluripotent stem cells are beginning to enter Phase 1 clinical trials (Piao et al., 2021; Takahashi, 2020), and there is a growing need for unbiased methods for maintaining and expanding high quality hPSCs. Currently, culture of hPSCs is a labor-intensive process requiring highly skilled operators. Human judgement is currently required for the most fundamental tasks: counting the number of cells in a culture to optimize timing and concentrations for passaging the cells and assessing the quality of cultures for contamination with cellular morphologies associated with unwanted differentiation. Artificial intelligence (AI) offers the speed and flexibility needed to bridge the gap between a human-dependent process and industrial-scale automation. We have designed an accurate method for hPSC visualization that ignores microscopic artifacts including particulate on the bottom of the microwell plate, well edge fluorescence, bubbles, and variation in focal planes. Our approach intentionally uses standard tissue culture plates and does not require augmentation of bright field images prior to training.

We tested three AI tasks for determining clone quality: distinguishing between clones, classifying images as undifferentiated or differentiated, and semantic segmentation. All were able to generally rank the hPSC clones based on quality that correlates with results from classical pluripotency assays, but we found that because spatial information is also obtained, semantic segmentation was a superior method for conducting hPSC quality assessments. While there are two examples of image-based classification of hPSCs to evaluate the presence of differentiated cells (Kusumoto *et al*., 2018; Piotrowski et al., 2021), neither is capable of assigning classifications at single pixel resolution, a clear strength of using AI semantic segmentation.

Machine learning is dependent on the quality of the input data and high importance is placed on having a sufficiently balanced and representative dataset. In our study, the AI was tolerant of image area sampled to determine the cell density, but it failed to generate accurate density maps when the image resolution was degraded due to the lack of contrast within a field of view. We found that preset hyperparameters generated convincing results and have not fully explored the hyperparameters associated with each of the neural networks tested.

Besides the basic tasks of cell counting and discovery of differentiated contaminants, the AI-generated density map contains information that would allow additional analyses. For example, it could be used for mapping cell positions, detecting confluency, and measuring inter-nuclear distances. The method we tested can be easily embedded into an automated process capable of a scale and throughput to meet the demands of automated hPSC culturing (Elanzew et al., 2020). The cell counting, and quality assessment methods can be adapted to a variety of hPSC lines and microscopes through training of new models or transfer learning with as little as a single 96-well plate of hPSCs. We also envision the method being used as a means for establishing standards when training individuals to conduct hPSC tissue culture work. With cell type- or microscopy-specific training data sets, we expect these methodologies to be expandable to assess other cell types, such as intermediate progenitors and differentiated derivatives of hPSCs. In that capacity, the AI methods may provide rapid quality control assessments for cells being cultured for use as cell replacement therapies, augmenting existing validation methods like gene expression profiling, flow cytometry, and immunocytochemical analyses.

## Materials and Methods

### Tissue culture

The hPSC line was obtained from Life Technologies (Thermo) and maintained between passages 25 to 45. The cell line is an episomal reprogrammed line derived from CD34+ hematopoietic somatic cells. hPSCs were fed daily using mTeSR1, passaged using ReLeSR, and attached to hESC-qualified Matrigel (Corning) coated 10cm and 96 well tissue culture plates. hPSCs were evaluated for pluripotency by flow cytometry, karyotypical abnormalities, and mycoplasma to control for quality of cultures.

The reprogrammed hPSC clones were derived from CD34+ cord blood cells (Stemcell Technologies). The reprogramming was done using the CytoTune-iPS 2.0 Sendai Reprogramming kit (Invitrogen) and following the instruction manual. Once reprogramming was completed the clones were fed daily using mTeSR1, passaged using ReLeSR, and attached to hESC-qualified Matrigel (Corning) coated 6 well tissue culture plates.

### Cell plating and staining for training dataset

The hPSC line was dissociated from a 10cm dish using ReLeSR and plated at equal densities on standard flat bottom 96 well microplates (Corning) that were coated with hESC-qualified matrigel. Plates were fixed on subsequent days in one day intervals. All plates were fixed with a final concentration of 3.7% formaldehyde for 20 minutes by adding an equal volume to the media already in the well of 7.4% formaldehyde. To stain the nuclei of the cells a staining solution was made by diluting Hoechst 33342 (Molecular Probes) to 1:5000 in PBS and incubating in the dark for 15 minutes at room temperature. After the incubation the staining solution was removed and the cells were washed three times with PBS and a sufficient volume (∼200 μL) of PBS was added to the wells for imaging.

### Imaging and acquisition settings

Images were acquired with the Celigo Imaging Cytometer (Nexcelom Bioscience). The illumination for brightfield is a 1 LED-based enhanced brightfield imaging channel with uniform well illumination. There are also 4 LED-based fluorescent channels. A large chip CCD camera along with galvanometric mirrors and an F-theta lens are used to acquire the images at a 1 μLm/pixel resolution. All images are at 10x magnification.

Training plates for the cell counting and density map model were imaged in both brightfield and blue channel. All other plates were imaged in brightfield only. Acquisition settings: Brightfield 50 ms exposure; Hoechst 250 ms exposure, excitation 377/50, emission 470/22

### Cell Counting & Density Map Model Training

The model was trained using Amazon SageMaker. An ml.p3.8xlarge instance was used for training. The custom model and training script from Sendagi, et al. (Sindagi and Patel, 2017) was packaged into a docker image according to SageMaker specifications. All hyperparameters used during training were kept the same as from Sendagi, et al. (Sindagi and Patel, 2017). Training lasted approximately 4 hours.

The training dataset was assembled by randomly selecting 3000 1958×1958 images from a larger dataset of 4608 images acquired from three 96 wells plates. Each image was then reduced in size to 256×256 by taking a random crop from the image. The 3000 images were then manually sorted to remove images that were out of focus or otherwise had defects preventing the nuclei segmentation algorithm from working properly. After manually sorting through the 3000 images we were left with 2375 good images to use for training. We then took the fluorescence channel from each image and ran it through a standard segmentation algorithm to find the nuclei center points. Those center points were then used to create the ground truth density map as described in (Sindagi and Patel, 2017). The training dataset was then further split into a training and validation dataset, with 80% of the data used for training and 20% of the data used for validation during training.

### Automated prototype platform

We created an automated platform to monitor hPSC growth by integrating a robotic arm (Precise Automation), Cytomat automated incubator (Thermo Scientific), Celigo Imaging Cytometer (Nexcelom) and Overlord laboratory automation and control software (Peak Analysis & Automation).

### Automated Runs to Confluency

The human hPSC line was dissociated from a 10cm dish using reLeSR and plated at equal densities on 96 well microplates that were coated with hESC-qualified matrigel. After plating the 96 well microplates were loaded into the Cytomat automated incubator in our prototype automated system. Using Overlord automation software the plates were set to image all plates and upload those images to an AWS S3 bucket every <u>12</u> hours. Images were run through the model on AWS. We used the model output of cell counts, heat maps and growth curves to track cell growth. Plates were maintained until cells grew to confluency.

### Continuous Run

The hPSC line was dissociated from a 10cm dish using reLeSR and plated at <u>four</u> different densities on each of the 96 well microplates that were coated with hESC-qualified matrigel. After plating the 96 well microplates were loaded into the <u>Cytomat</u> automated incubator in our prototype automated system. Using Overlord automation software the plates were set to image all plates and upload those images to AWS every 12 hours. We used the model output to determine when plates were ready to split and the split ratio to be used to equilibrate cell densities across each microplate.

### hPSC Quality Classification Model Training

These models were all trained using MATLAB 2020b running in an AWS EC2 p3.2xlarge instance. To create a training set for clone identification, 1,000 random images from each clone were selected for a total of 7,000 images. Of these, 60% were used for training, 20% were used for validation during training, and 20% were used for testing and evaluating the trained model. The pretrained model used for clone identification with transfer learning was densenet201. The final model had a validation accuracy of 79.36%.

| Hyperparameter | Value |
| --- | --- |
| Solver | SDGM |
| Momentum | 0.9 |
| Initial Learning Rate | 0.001 |
| Learn Rate Schedule | Piecewise |
| Learn Rate Drop Factor | 0.1 |
| Learn Rate Drop Period | 10 |
| L2 Regularization | 1.0000e-04 |
| Gradient Threshold Method | L2norm |
| Gradient Threshold | Inf |
| Max Epochs | 10 |
| Mini Batch Size | 100 |
| Verbose | 1 |
| Verbose Frequency | 50 |
| Validation Frequency | 20 |
| Validation Patience | 5 |
| Shuffle | Every epoch |
| Execution Environment | Single GPU |

To create a training set for determining clone quality, images from several hPSC clones and the hPSC line from Life Technologies were used and separated into hPSC and non-hPSC classes. The original images were acquired in 6 well plates and were size 1958×1958, and were tiled into four 979×979 images to use for training. From the complete set of clone images 1200 were selected for each class for a total of 2400 images. Of these, 60% were used for training, 20% were used for validation during training, and 20% were used for testing and evaluating the trained model. The pretrained model used for determining clone quality with transfer learning was resnet101. The final model had a validation accuracy of 95.15%.

| **Hyperparameter** | **Value** |
| --- | --- |
| Solver | SDGM |
| Momentum | 0.9 |
| Initial Learning Rate | 0.001 |
| Learn Rate Schedule | Piecewise |
| Learn Rate Drop Factor | 0.1 |
| Learn Rate Drop Period | 10 |
| L2 Regularization | 1.0000e-04 |
| Gradient Threshold Method | L2norm |
| Gradient Threshold | Inf |
| Max Epochs | 10 |
| Mini Batch Size | 100 |
| Verbose | 1 |
| Verbose Frequency | 50 |
| Validation Frequency | 10 |
| Validation Patience | 5 |
| Shuffle | Every epoch |
| Execution Environment | Single GPU |

### hPSC Semantic Segmentation Model Training

These models were all trained using MATLAB 2020b running in an AWS EC2 p3.2xlarge instance. To create a training set to segment hPSC, non-hPSC and background in images, 32 random images of each hPSC clone were selected for a total of 224 images. Of these, 60% were used for training, 20% were used for validation during training, and 20% were used for testing and evaluating the trained model. The pixel labels were created using MATLAB Image Labeler to label pixels as hPSC, non-hPSC or background. The semantic segmentation network used to train this model was Deeplabv3+ and the base pretrained network was resnet50. The final model had a validation accuracy of 95.99%, a weighted intersection over union (IoU) score of 0.94, and a mean boundary F1 (BF) score of 0.792. The IoU and BF scores are calculated on the training dataset. The IoU is the ratio of correctly classified pixels to the total number of ground truth and predicted pixels in that class. The BF score shows how well the predicted boundary of each class aligns with the true boundary.

| **Hyperparameter** | **Value** |
| --- | --- |
| Solver | SDGM |
| Momentum | 0.9 |
| Initial Learning Rate | 1.0000e-05 |
| Learn Rate Schedule | Piecewise |
| Learn Rate Drop Factor | 0.1 |
| Learn Rate Drop Period | 3 |
| L2 Regularization | 1.0000e-04 |
| Gradient Threshold Method | L2norm |
| Gradient Threshold | Inf |
| Max Epochs | 7 |
| Mini Batch Size | 5 |
| Verbose | 0 |
| Verbose Frequency | 50 |
| Validation Frequency | 15 |
| Validation Patience | 5 |
| Shuffle | Every epoch |
| Execution Environment | Single GPU |

### Reagent/Resource Source Identifier

Human Episomal iPSC Line Gibco by Life Technologies A18945

mTeSR1 Stemcell Technologies 85850

ReLeSR Stemcell Technologies 05873

DMEM/F-12 Gibco 10565018

hESC-qualified Matrigel Corning 354277

Clear 96 well microplate Corning 353872

Clear 6 well microplate Corning 353846

10cm Dish Nunc 150464

Formaldehyde Sigma Aldrich 252549

DPBS Gibco by Life Technologies 14190144

Hoescht 33342 Molecular Probes by Life Technologies H3570

CytoTune-iPS 2.0 Sendai Reprogramming Kit Invitrogen A16517

Human CD34+ Cord Blood Cells Stemcell Technologies 70008

## Acknowlegements

We would like to thank Roy Williams for help in imputing PluriTest results of the the mRNA bulk sequencing data. This research was funded in part by a California Institute of Regenerative Medicine grant, EDUC2-08394.

---

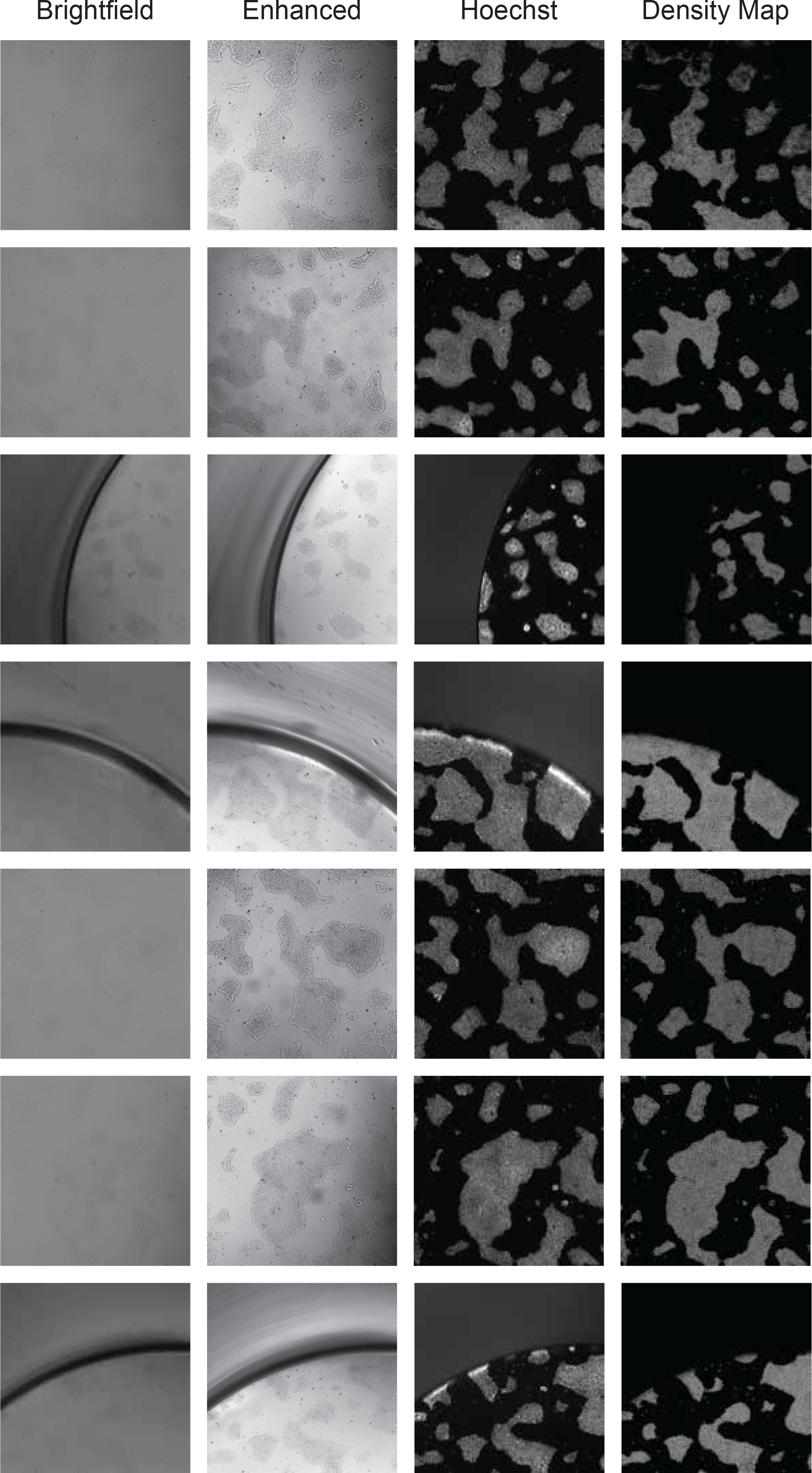
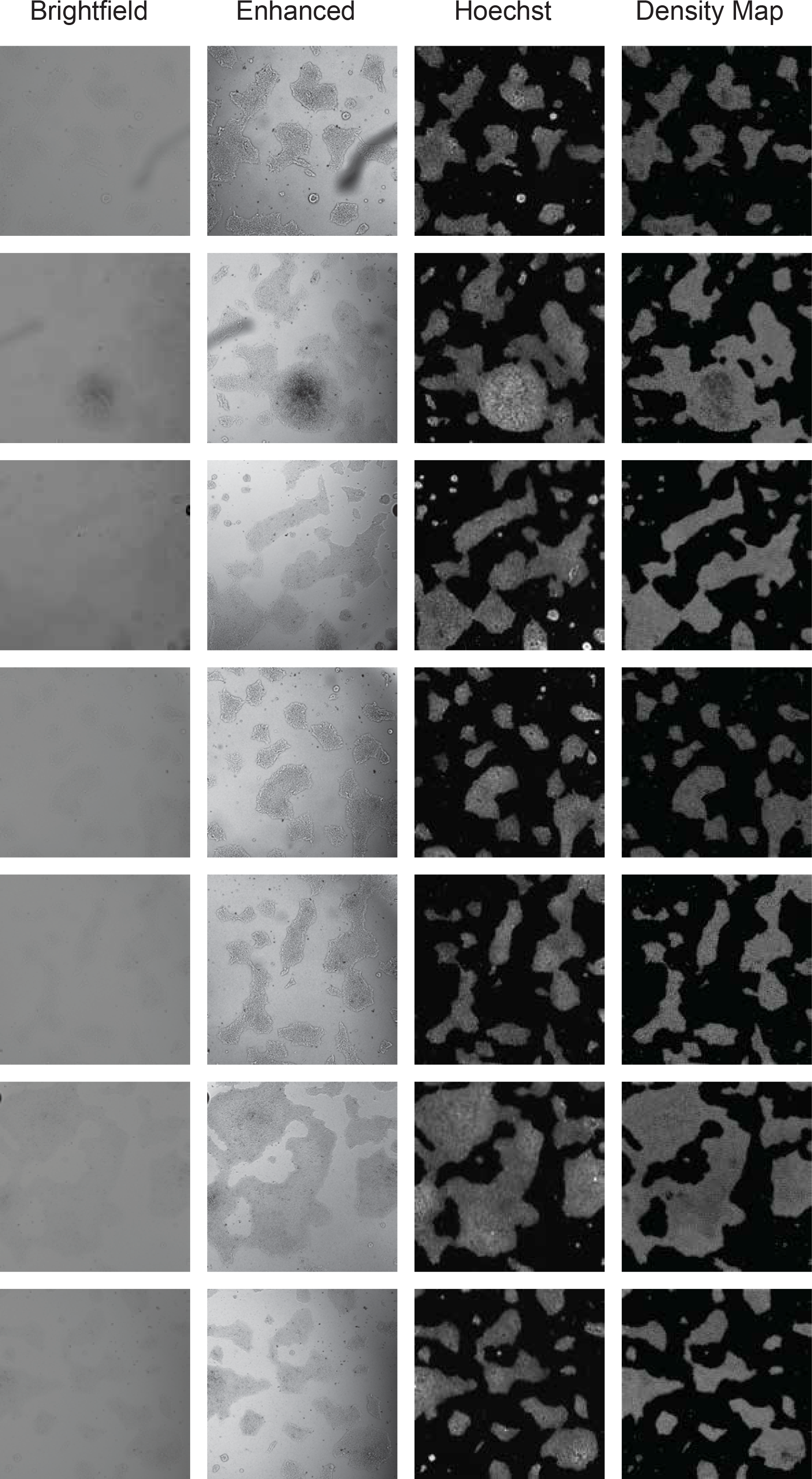
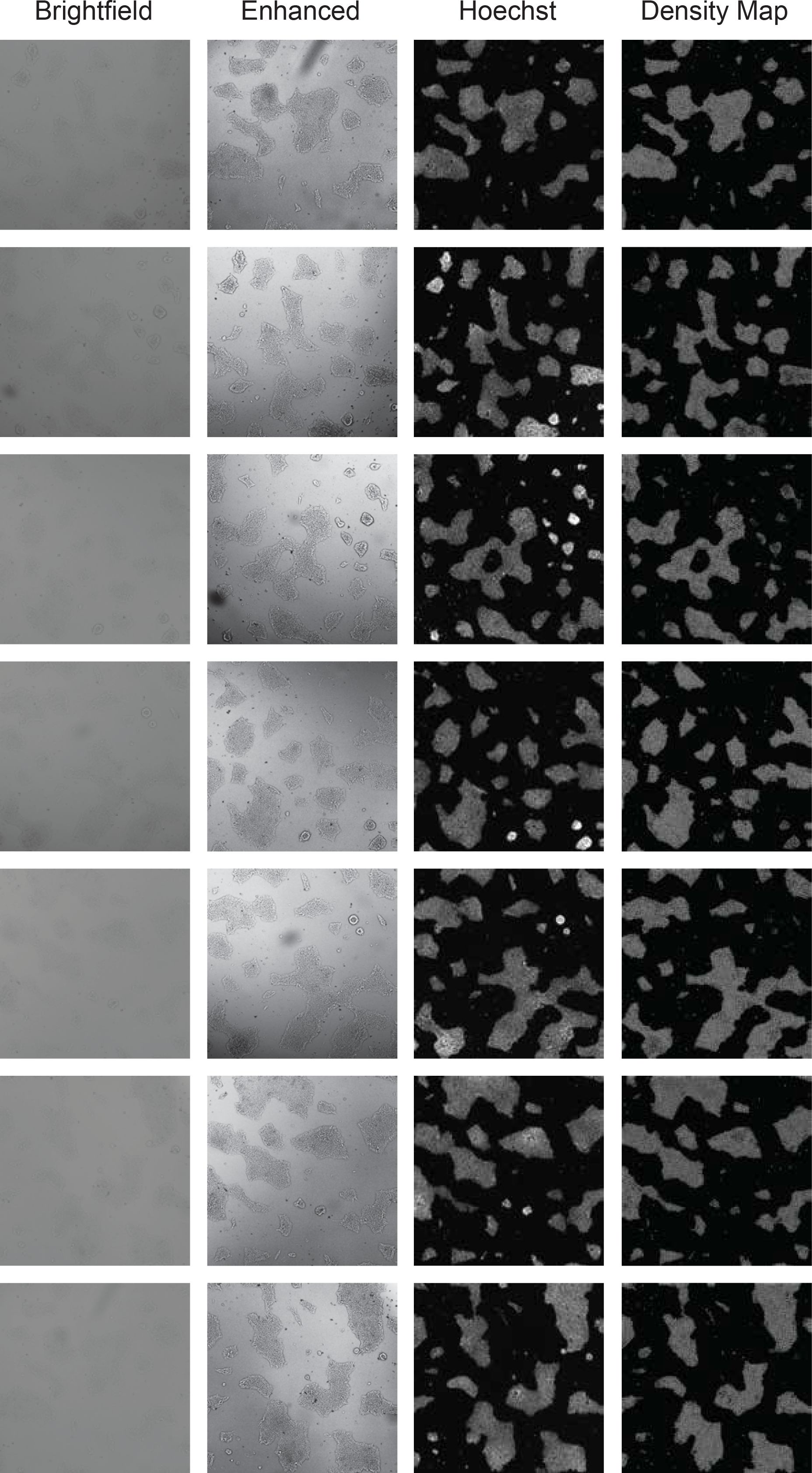
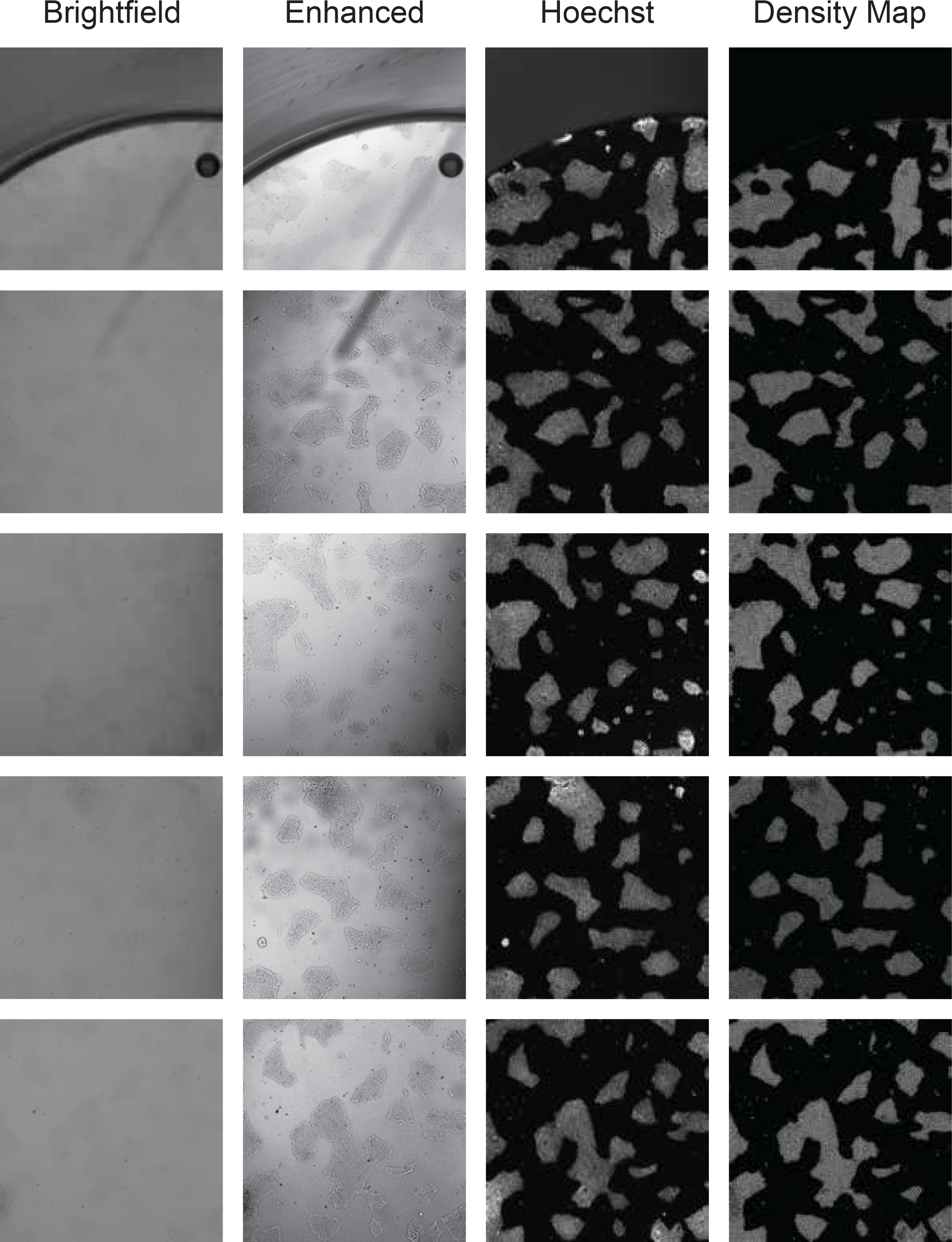

**Supplemental Table 1.**
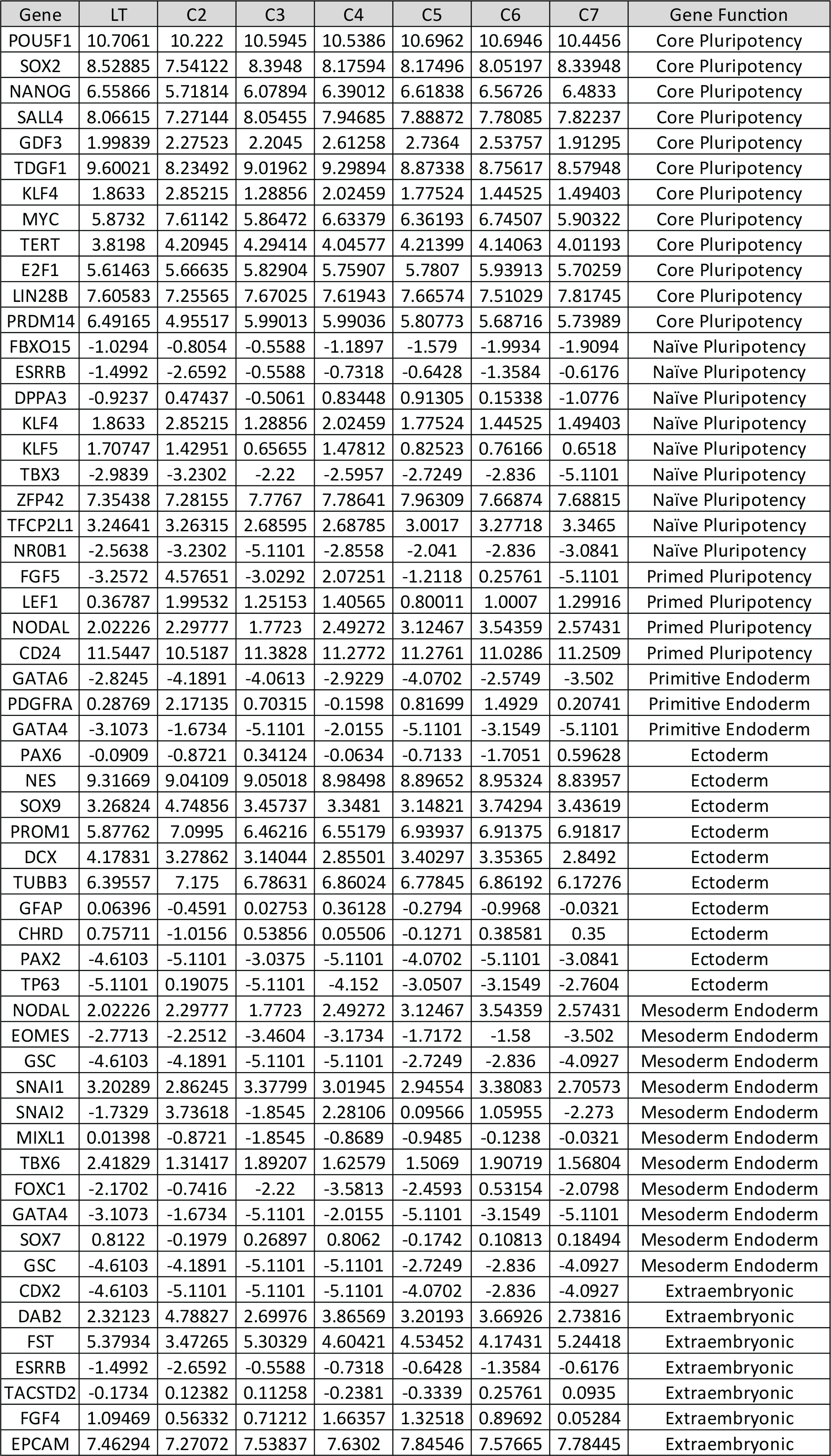
Gene expression data for the iPSC lines focusing on key genes for pluripotency and differentiation.

**Supplemental Table 2.**
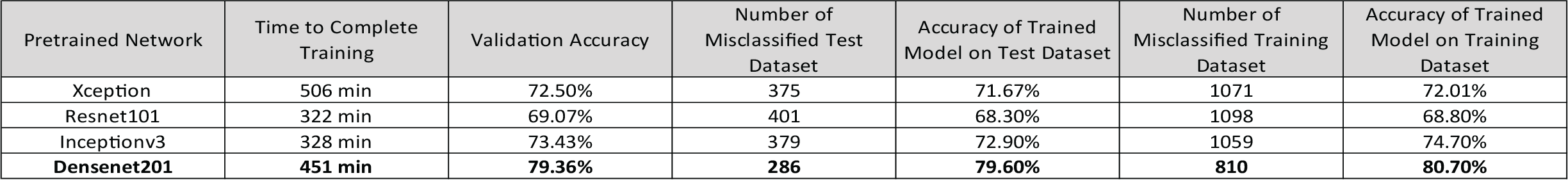
Pretrained networks tested for transfer learning of classification networks. To train a model to distinguish between the different iPSC clones, Densenet201 was selected because it resulted in the highest validation and test set accuracy.

| Pretrained Network | Time to Complete Training | Validation Accuracy | Number of Misclassified Test Dataset | Accuracy of Trained Model on Test Dataset | Number of Misclassified Training Dataset | Accuracy of Trained Model on Training Dataset |
| --- | --- | --- | --- | --- | --- | --- |
| Xception | 506 min | 72.50% | 375 | 71.67% | 1071 | 72.01% |
| Resnet101 | 322 min | 69.07% | 401 | 68.30% | 1098 | 68.80% |
| Inceptionv3 | 328 min | 73.43% | 379 | 72.90% | 1059 | 74.70% |
| **Densenet201** | **451 min** | **79.36%** | **286** | **79.60%** | **810** | **80.70%** |

**Supplemental Table 3.** Pretrained networks tested for transfer learning of classification networks. To train a model to distinguish between pluripotent and differentiated iPSC images, Resnet101 was selected because of the relatively short training time and high accuracy.

| Pretrained Network | Time to Complete Training | Validation Accuracy | Number of Misclassified Test Dataset | Accuracy of Trained Model on Test Dataset | Number of Misclassified Training Dataset | Accuracy of Trained Model on Training Dataset |
| --- | --- | --- | --- | --- | --- | --- |
| Xception | 124 min | 92.72% | 23 | 94.40% | 100 | 91.90% |
| Inceptionv3 | 83 min | 92.96% | 22 | 94.70% | 45 | 96.40% |
| Densenet201 | 135 min | 96.36% | 13 | 96.80% | 6 | 99.50% |
| **Resnet101** | **69 min** | **95.87%** | **7** | **98.30%** | **5** | **99.60%** |

**Supplemental Table 4.** Pretrained networks tested as base networks for DeeplabV3+ for transfer learning of semantic segmentation networks. To train a model to segment individual pixels in images as iPSC, non iPSC or background Resnet50 was selected because of the relatively short training time and high Accuracy, IoU and BF1 scores.

| Base Network | Time to Complete Training | Validation Accuracy | Global Accuracy of Trained Model on Test Dataset | Weighted IoU Score of Trained Model on Test Dataset | Mean BF Score of Trained Model on Test Dataset | iPSC pixel prediction Accuracy of Trained Model on Test Dataset | Non iPSC pixel prediction Accuracy of Trained Model on Test Dataset | Global Accuracy of Trained Model on Training Dataset | Weighted IoU Score of Trained Model on Training Dataset | Mean BF Score of Trained Model on Training Dataset | iPSC pixel prediction Accuracy of Trained Model on Training Dataset | Non iPSC pixel prediction Accuracy of Trained Model on Training Dataset |
| --- | --- | --- | --- | --- | --- | --- | --- | --- | --- | --- | --- | --- |
| Mobilenetv2 | 10 min | 94.86% | 0.941 | 0.863 | 0.689 | 0.96 | 0.85 | 0.95 | 0.86 | 0.708 | 0.96 | 0.82 |
| Xception | 64 min | 92.71% | 0.918 | 0.824 | 0.647 | 0.91 | 0.89 | 0.939 | 0.848 | 0.667 | 0.92 | 0.89 |
| Resnet18 | 10 min | 96.05% | 0.946 | 0.9 | 0.735 | 0.94 | 0.91 | 0.968 | 0.94 | 0.776 | 0.97 | 0.95 |
| **Resnet50** | **11 min** | **95.99%** | **0.965** | **0.934** | **0.768** | **0.98** | **0.86** | **0.968** | **0.94** | **0.792** | **0.97** | **0.95** |

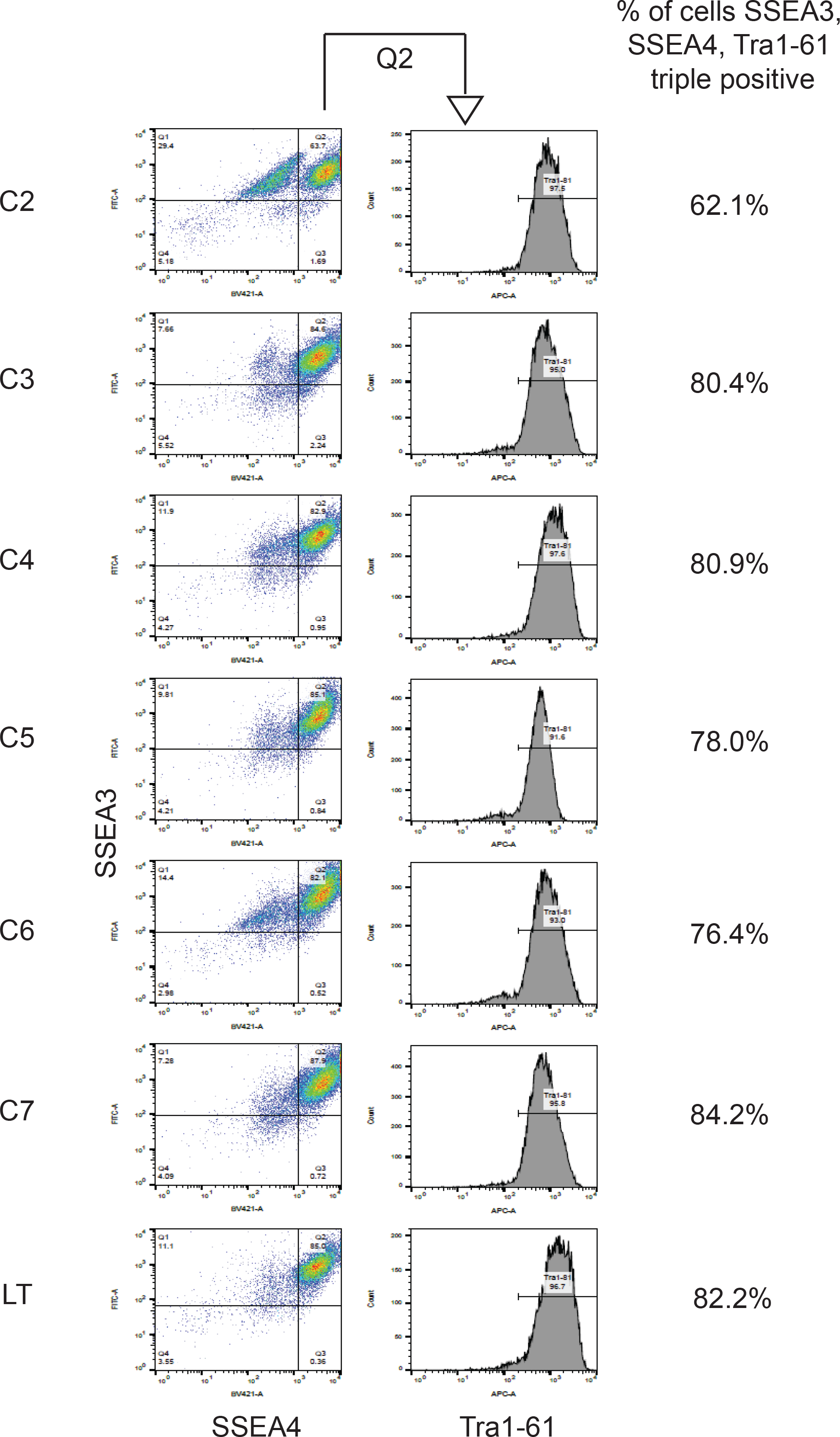

